# Tetrapeptide caspase inhibitors exhibit protease- and substrate-dependent off-target specificity

**DOI:** 10.64898/2026.11.13.766510

**Authors:** Christopher M. Bourne, Jacob D. Liebovitz, Jordi C.J. Hintzen, Marzieh Daniali, Kithya Borey, Haley T. Goodrow, Elizabeth Mitroka, Bohdana M. Discher, George M. Burslem, Peter J. Gaskill, Cornelius Y. Taabazuing

## Abstract

Peptide-based caspase inhibitors are widely used to define the functions of individual caspases and were developed from tetrapeptide substrate preferences identified using positional scanning peptide libraries. Their specificity is typically validated using fluorogenic tetrapeptide substrates, yet whether these assays accurately predict inhibition of endogenous protein substrates has not been systematically examined. Here, we demonstrate that the widely used executioner caspase inhibitor zDEVD-FMK directly inhibits caspase-1 (CASP1)-mediated processing of interleukin-1β (IL-1β) and interleukin-18 (IL-18) despite CASP1 exhibiting minimal activity toward the DEVD fluorogenic substrate. Unexpectedly, zDEVD-FMK selectively inhibited CASP1 but not the closely related inflammatory caspases CASP4 or CASP5. Conversely, the CASP1 inhibitor Ac-YVAD-CMK directly inhibited CASP3-mediated protein substrate cleavage. We further found that endogenous substrates displayed markedly different sensitivities to inhibition, with IL-1β and IL-18 processing consistently inhibited more efficiently than gasdermin D cleavage, while fluorogenic CASP3/7 reporter cleavage substantially overestimated inhibition of endogenous apoptotic substrates. During inflammasome activation and *Salmonella* infection, DEVD- and YVAD-based inhibitors failed to distinguish apoptotic executioner caspase activity from inflammatory caspase signaling. Together, our findings demonstrate that fluorogenic tetrapeptide assays do not faithfully predict inhibition of endogenous protein substrates and identify protease- and substrate-dependent specificity as fundamental features of peptide-based caspase inhibitors.

## Introduction

Programmed cell death is essential for development, tissue homeostasis, and host defense against infection and cancer^1–5^. Caspases are cysteine proteases that initiate and execute distinct cell death programs with diverse immunological consequences^6,7^. Apoptosis is a non-lytic form of programmed cell death that is generally considered immunologically silent and is initiated by caspase-9 through the intrinsic pathway or caspase-8 through the extrinsic pathway^7,8^. The intrinsic pathway activates caspase-9 through the apoptosome, whereas the extrinsic pathway activates caspase-8 through the death-inducing signaling complex (DISC)^7,9,10^. Both pathways converge on the executioner caspases-3 and -7, which dismantle the cell by cleaving proteins required for DNA repair, cytoskeletal integrity, vesicle trafficking, and gene expression^11^.

In contrast, pyroptosis is a lytic, pro-inflammatory form of programmed cell death triggered by host recognition of damage-associated molecular patterns (DAMPs) and pathogen-associated molecular patterns (PAMPs)^6,12^. Recognition of these danger signals induces assembly of inflammasomes, large macromolecular signaling complexes that promote oligomerization and autoproteolytic activation of the inflammatory caspases-1, -4, and -5 in humans and -11 in mice^13–17^. In the canonical pathway, inflammasome sensor proteins recruit the adaptor apoptosis-associated speck-like protein containing a CARD (ASC) and CASP1 to form the canonical inflammasome complex^18^. Activated CASP1 cleaves the pore-forming protein gasdermin D (GSDMD), whose N-terminal fragment forms membrane pores that drive osmotic imbalance, cell swelling, and lytic cell death^19–21^. CASP1 also processes pro-IL-1β and pro-IL-18 into their mature forms, which are released through GSDMD pores to amplify inflammation^22–25^. In the non-canonical pathway, cytosolic lipopolysaccharide (LPS) directly binds CASP4 or CASP5, triggering their activation and subsequent GSDMD cleavage to induce pyroptosis ^26,27^. CASP4 and CASP5 also process IL-18 to further promote inflammatory responses^28–30^.

Although apoptosis and pyroptosis were historically viewed as distinct cell death programs, growing evidence demonstrates extensive bidirectional crosstalk between these pathways^14,31–34^. CASP1 can activate both initiator and executioner caspases to induce apoptosis^31,34–36^. Likewise, we recently demonstrated that CASP4 and CASP5 directly cleave and activate CASP3 and CASP7, establishing a direct link between non-canonical inflammasome activation and apoptotic executioners^37^. Conversely, CASP3 regulates inflammatory signaling by cleaving and inactivating GSDMD, IL-1β, and IL-18, but CASP3 can also convert apoptosis into pyroptosis through cleavage of GSDME^31,38–41^. Notably, physiologic stimuli such as bacterial infection frequently activate apoptotic and pyroptotic pathways simultaneously, making it difficult to assign cellular outcomes, including cytokine maturation, membrane permeabilization, and cell death, to either pathway in isolation. Dissecting the relative contributions of these pathways therefore requires pharmacological tools capable of selectively inhibiting individual caspases without perturbing the others.

The development of such tools has been guided by the observation that caspases recognize tetrapeptide motifs immediately N-terminal to the cleavage site^42,43^. Positional scanning combinatorial peptide libraries identified the optimal recognition sequence for individual caspases, including the Asp-Glu-Val-Asp (DEVD) motif preferentially recognized by the executioner caspases-3 and -7^44,45^. This sequence formed the basis for irreversible peptide inhibitors containing electrophilic warheads that covalently modify the catalytic cysteine and remain among the most widely used tools for studying apoptosis. However, substantial cross-reactivity has been reported among inhibitors targeting apoptotic initiator and executioner caspases^46^, likely reflecting the high degree of structural conservation within the caspase family. Whether DEVD-based inhibitors can distinguish apoptotic executioner caspases from the structurally similar but functionally distinct inflammatory caspases has not been systematically examined. While using DEVD-based inhibitors to block executioner caspases, we unexpectedly observed inhibition of inflammatory caspase substrate processing, prompting us to investigate their specificity.

Here, we demonstrate that DEVD-based inhibitors directly inhibit CASP1, but not CASP4 or CASP5, in both biochemical assays and intact cells. Unexpectedly, DEVD inhibitors preferentially inhibit CASP1-mediated processing of IL-1β and IL-18 over GSDMD in immortalized cell lines, but inhibit GSDMD over IL-1β in primary cells, revealing substrate-dependent inhibition of endogenous CASP1 cleavage. Reciprocally, the canonical CASP1 inhibitor YVAD directly inhibits CASP3 activity. Together, our findings demonstrate that tetrapeptide substrate assays do not faithfully predict endogenous protein substrate inhibition and that DEVD- and YVAD-based inhibitors cannot reliably distinguish apoptotic from inflammatory caspase signaling when both pathways are activated. These findings establish substrate-dependent inhibition as a previously unrecognized determinant of caspase inhibitor specificity and underscore the need for next-generation caspase inhibitors with improved protease and substrate selectivity.

## Results

### DEVD-based inhibitors selectively inhibit CASP1 and differentially block CASP1 substrates *in vitro*

Fluorogenic tetrapeptide substrates are routinely used to evaluate caspase activity during inhibitor development. However, cleavage of short synthetic peptides does not necessarily predict inhibition of endogenous protein substrates, whose recognition is influenced by structural context and exosite interactions. We therefore asked whether the widely used executioner caspase inhibitor zDEVD-FMK directly inhibits inflammatory caspase cleavage of protein substrates.

To assess this, recombinant CASP1, CASP4, or CASP5 were incubated with a four-fold dilution series of zDEVD-FMK (starting at 10 μM) before the addition of THP-1 macrophage lysates, and substrate cleavage was assessed by immunoblotting. Unless otherwise stated, all recombinant caspases were assayed at 1 activity unit/μL. Unexpectedly, zDEVD-FMK potently inhibited CASP1-mediated processing of IL-18 and IL-1β, with inhibition evident at concentrations as low as 0.16 μM (Figure 1A). In contrast, cleavage of the CASP1 substrate GSDMD was not inhibited until 10 μM, indicating that individual CASP1 substrates differ markedly in their sensitivity to inhibition (Figure 1A). Whereas CASP1-mediated cytokine processing was potently inhibited, zDEVD-FMK had no detectable effect on CASP4 or CASP5-mediated cleavage of IL-18 or GSDMD (Figure 1B,C). As expected, zDEVD-FMK inhibited PARP cleavage by the executioner caspases, completely blocking CASP3 activity at 0.63 μM and CASP7 activity at 10 μM (Figure 1D,E), consistent with previous reports demonstrating that PARP is a preferred CASP7 substrate^47^. Thus, CASP1-mediated cytokine processing was inhibited over the same concentration range required to inhibit CASP3 activity.

**Figure 1.**
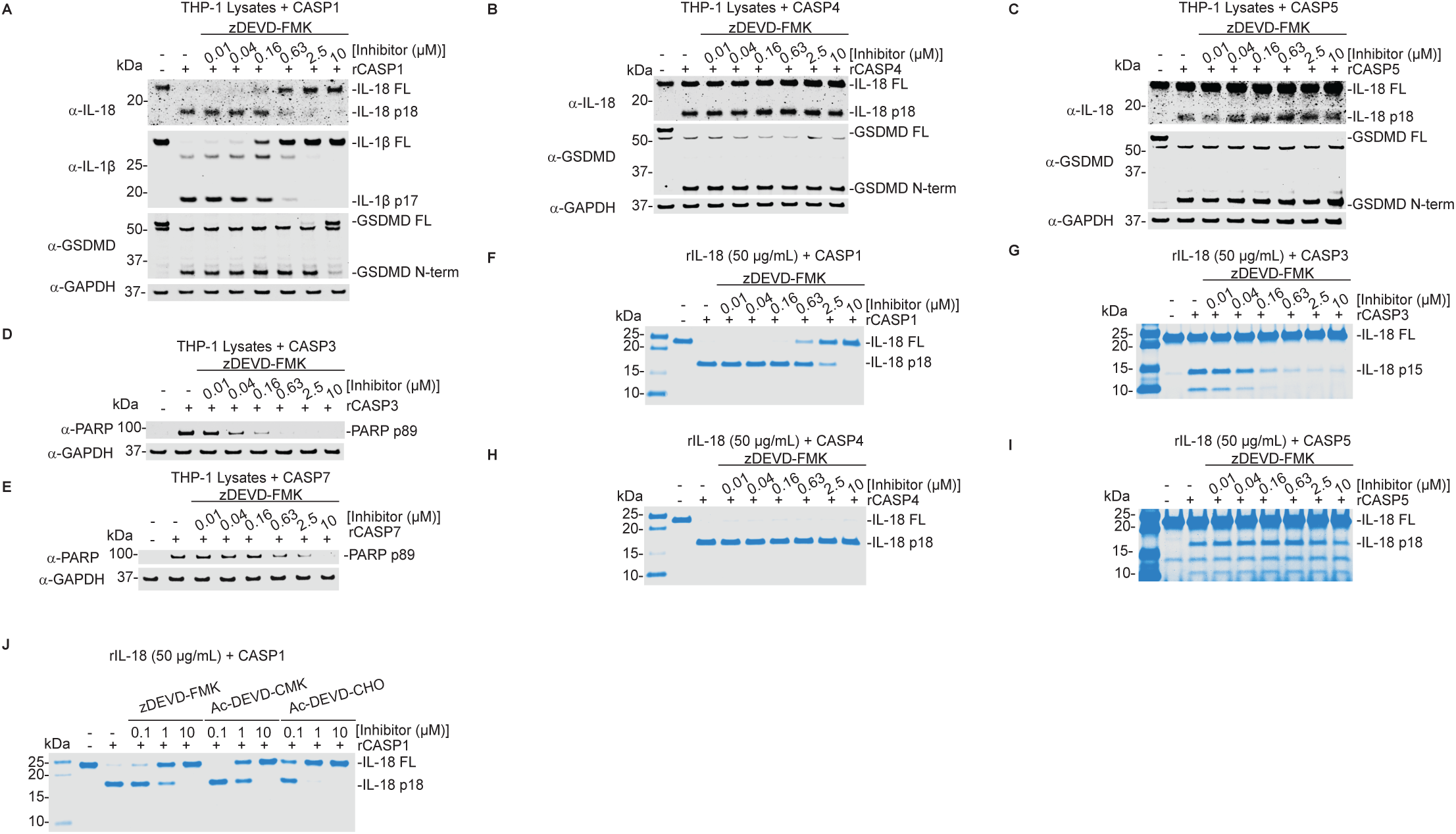
DEVD-based inhibitors selectively inhibit CASP1 and reveal substrate-dependent inhibition *in vitro*. (A-E) Recombinant active caspases (1 activity unit/μL) were preincubated with four-fold dilution series of zDEVD-FMK (starting at 10 μM) before the addition of THP-1 macrophage lysates. Cleavage of IL-18, IL-1β, and GSDMD by CASP1 (A), IL-18 and GSDMD by CASP4 (B), IL-18 and GSDMD by CASP5 (C), PARP by CASP3 (D), and PARP by CASP7 (E) was assessed by immunoblotting. (F-I) Recombinant active caspases were preincubated with zDEVD-FMK before the addition of recombinant IL-18 (50 μg/mL). IL-18 cleavage by CASP1 (F), CASP3 (G), CASP4 (H), and CASP5 (I) was analyzed by Coomassie staining. (J) Recombinant CASP1 was preincubated with zDEVD-FMK, Ac-DEVD-CMK, or Ac-DEVD-CHO (0.1, 1, or 10 μM) before the addition of recombinant IL-18 (50 μg/mL), and IL-18 cleavage was analyzed by Coomassie staining. Data are representative of at least three independent experiments.

Because THP-1 lysates contain numerous competing caspase substrates that could influence the apparent potency of zDEVD-FMK, we next examined cleavage of purified recombinant IL-18. IL-18 provides a common substrate because it is cleaved by inflammatory caspases at D36 and by CASP3 at D71 and D76, generating an inactive fragment^40^. Recombinant CASP1, CASP3, CASP4, or CASP5 were incubated with recombinant IL-18 in the presence of a four-fold dilution series of zDEVD-FMK, and cleavage products were analyzed by Coomassie staining. Consistent with the lysate-based assays, zDEVD-FMK inhibited CASP1-mediated IL-18 cleavage beginning at 0.63 μM (Figure 1F). CASP3-mediated IL-18 cleavage was substantially reduced at similar concentrations, although complete inhibition was not achieved within the concentration range tested (Figure 1G). In contrast, zDEVD-FMK did not inhibit CASP4-mediated IL-18 cleavage (Figure 1H) and only weakly inhibited CASP5 at the highest concentration tested (Figure 1I). These findings demonstrate that zDEVD-FMK directly inhibits CASP1-mediated protein substrate cleavage but exhibits little or no activity toward CASP4 or CASP5, indicating that DEVD-based inhibitors do not uniformly target inflammatory caspases.

The selective inhibition of CASP1 over the closely related inflammatory caspases CASP4 and CASP5 was unexpected. We therefore asked whether CASP1 inhibition was determined by the DEVD tetrapeptide sequence or by the N-terminal protecting group or C-terminal reactive warhead. Recombinant CASP1 was incubated with a ten-fold dilution series of zDEVD-FMK, Ac-DEVD-CMK, or Ac-DEVD-CHO before the addition of recombinant IL-18. Both zDEVD-FMK and Ac-DEVD-CMK partially inhibited IL-18 cleavage at 1 μM, whereas Ac-DEVD-CHO completely blocked cleavage at the same concentration (Figure 1J). These findings indicate that inhibition of CASP1 is an intrinsic property of DEVD-based inhibitors and is determined primarily by the DEVD recognition sequence rather than the N-terminal protecting group or reactive warhead. Together, these findings indicate that DEVD-based inhibitors selectively inhibit CASP1 but not the closely related inflammatory caspases CASP4 and CASP5. Furthermore, even within CASP1, endogenous substrates are not inhibited equally, with IL-1β and IL-18 processing being substantially more sensitive to inhibition than GSDMD cleavage.

### zDEVD-FMK covalently modifies the catalytic cysteine of CASP1 and CASP3 but not CASP4

Our biochemical assays demonstrated that zDEVD-FMK directly inhibits CASP1 and CASP3 but has little effect on the closely related inflammatory caspases CASP4 and CASP5. To investigate the molecular basis for this differential inhibition, we asked whether zDEVD-FMK covalently modifies the catalytic cysteine of CASP1 and CASP3 but not CASP4. We therefore performed liquid-chromatography tandem mass spectrometry on tryptic digests to identify and compare inhibitor-modified catalytic cysteine-containing peptides from recombinant CASP1, CASP3, and CASP4.

Each caspase was incubated with zDEVD-FMK or the pan-caspase inhibitor zVAD-FMK before proteolytic digestion and mass spectrometry analysis. To determine whether the inhibitors competed for the same catalytic residue, we also preincubated each caspase with zVAD-FMK for 30 min before adding zDEVD-FMK for an additional 30 min. Tandem mass spectrometry identified zVAD-FMK-modified catalytic cysteine peptides from all three caspases (Figure 2A,B). In contrast, zDEVD-FMK-modified peptides were detected for CASP1 and CASP3 but not CASP4 (Figure 2A,C). Preincubation with zVAD-FMK prevented detectable zDEVD-FMK modification of CASP1 and CASP3, consistent with competition for the catalytic cysteine (Figure 2A). No inhibitor modifications were detected outside the catalytic cysteine among the peptides analyzed. Altogether, these findings demonstrate that zDEVD-FMK covalently modifies the catalytic cysteine of CASP1 and CASP3, consistent with their sensitivity to inhibition. In contrast, the absence of detectable zDEVD-FMK modification of CASP4, despite its modification by zVAD-FMK, suggests that differential active-site engagement contributes to the distinct inhibitory profiles of inflammatory caspases.

**Figure 2.**
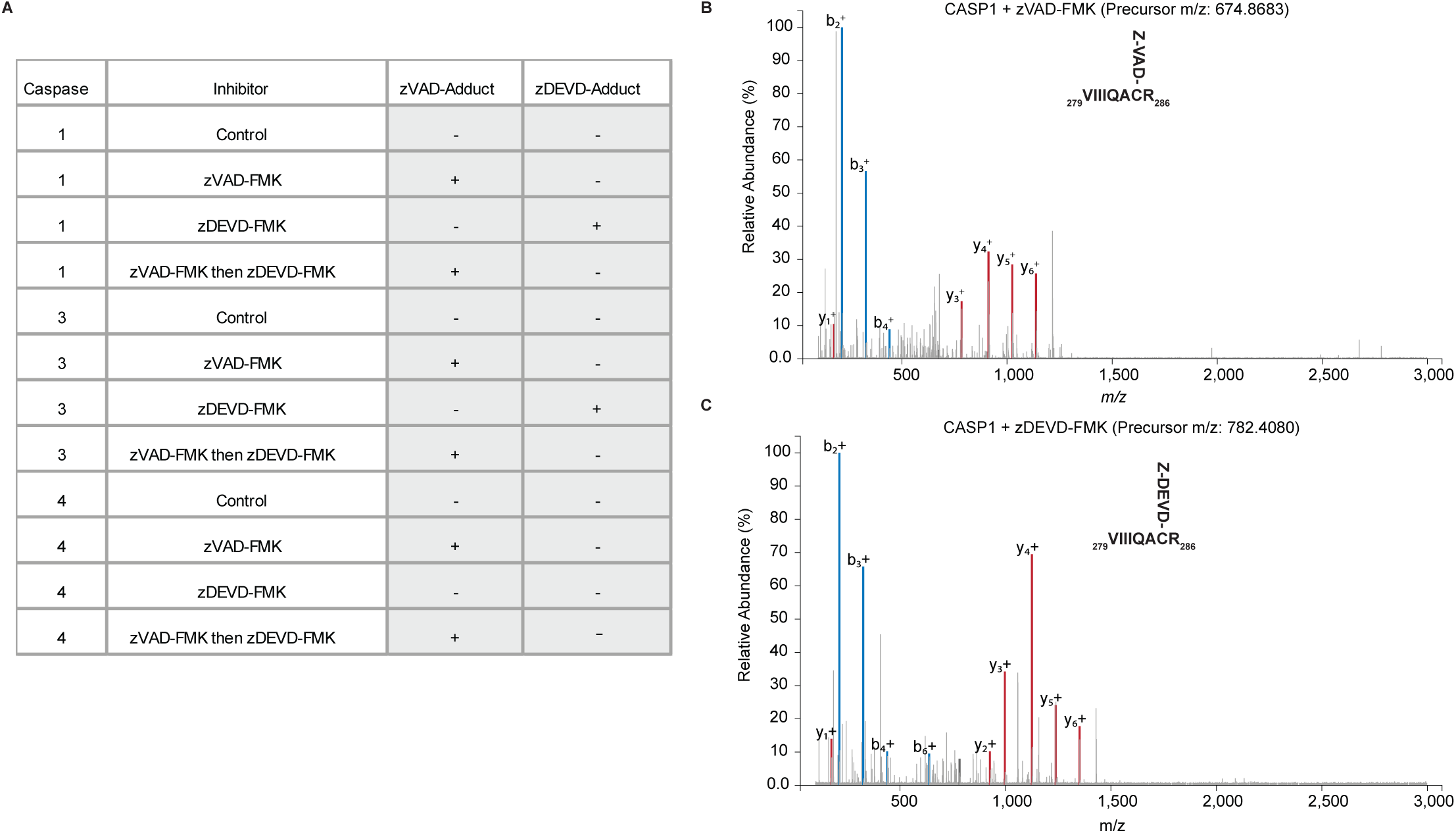
zDEVD-FMK covalently modifies the catalytic cysteine of CASP1 and CASP3 but not CASP4. LC-MS/MS analysis of inhibitor modification of the catalytic cysteine in recombinant CASP1, CASP3, and CASP4 following treatment with zVAD-FMK, zDEVD-FMK, or sequential treatment with both inhibitors. (A) Summary of detected inhibitor adducts on the catalytic cysteine of each caspase under the indicated conditions. (B,C) Representative MS/MS spectra of the CASP1 catalytic cysteine-containing peptide modified by zVAD-FMK at C285 (B) or zDEVD-FMK at C285 (C).

### Inhibition of fluorogenic probe cleavage does not predict inhibition of endogenous protein substrate cleavage

Because tetrapeptide-based caspase inhibitors were developed using positional scanning peptide libraries and fluorogenic substrates, we next asked whether inhibition of fluorogenic probe cleavage accurately predicts inhibition of endogenous protein substrates. Before addressing this question, we first confirmed that the recombinant caspases used throughout this study exhibited the expected tetrapeptide substrate preferences.

To do so, we performed Michaelis-Menten kinetic analyses using the fluorogenic probes Ac-WEHD-AMC and Ac-DEVD-AMC, the preferred substrates for CASP1 and CASP3, respectively. As expected, CASP1 efficiently cleaved Ac-WEHD-AMC but showed minimal activity toward Ac-DEVD-AMC (Figure 3A), exhibiting a 62-fold higher relative catalytic efficiency (K_cat_/K_m_) for Ac-WEHD-AMC (Figure 3C), consistent with previous reports (43). Conversely, CASP3 preferentially cleaved Ac-DEVD-AMC, with a 173.5-fold higher relative K_cat_/K_m_ than Ac-WEHD-AMC (Figure 3B,C). These data confirm that the recombinant enzymes display the expected tetrapeptide selectivity.

**Figure 3.**
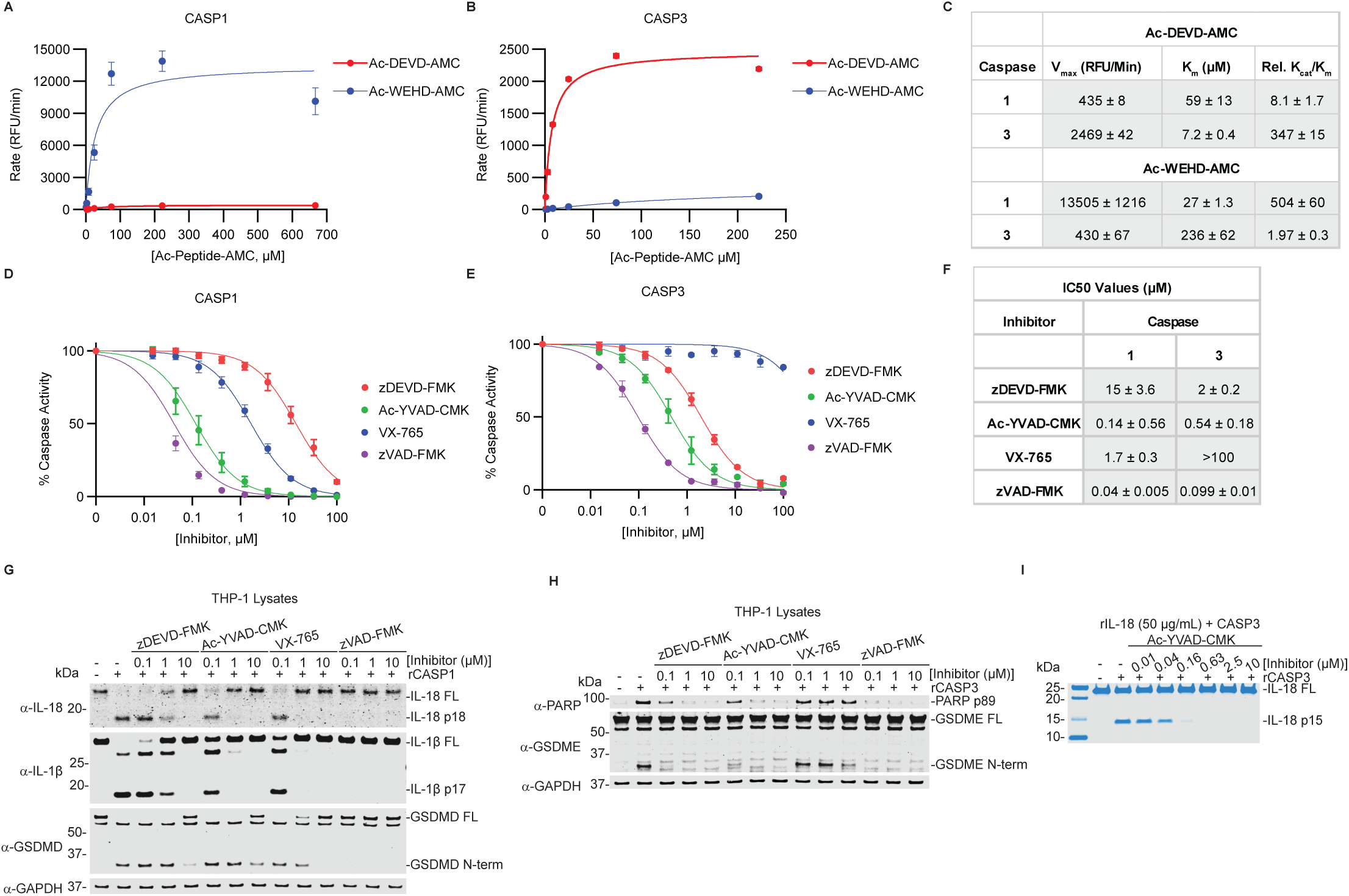
Inhibition of fluorogenic probe cleavage does not predict inhibition of endogenous protein substrate cleavage. (A,B) Michaelis-Menten kinetics of the preferred fluorogenic substrates for CASP1 (A) and CASP3 (B). (C) Table summarizing the V_max_, K_m_, and relative K_cat_/K_m_ values derived from panels (A) and (B). (D,E) Inhibition of fluorogenic substrate cleavage by recombinant CASP1 (D) or CASP3 (E) in the presence of zDEVD-FMK, Ac-YVAD-CMK, VX-765, or zVAD-FMK. (F) Calculated IC_50_ values for the fluorogenic probe assays shown in (D) and (E). (G,H) Immunoblot analysis of endogenous substrate cleavage by recombinant CASP1, showing IL-18, IL-1β, and GSDMD processing (G), or by recombinant CASP3 showing PARP and GSDME cleavage (H), following treatment with the indicated inhibitors. (I) Direct inhibition of recombinant CASP3-mediated IL-18 cleavage by Ac-YVAD-CMK analyzed by Coomassie staining. Data are presented as mean ± SEM from three independent replicates and are representative of at least three independent experiments.

Having confirmed the expected substrate preferences of CASP1 and CASP3, we next characterized the potency of each inhibitor using the preferred fluorogenic probe for each caspase. Recombinant CASP1 was incubated with zDEVD-FMK, Ac-YVAD-CMK, VX-765, or the pan-caspase inhibitor zVAD-FMK before the addition of saturating Ac-WEHD-AMC. Under these conditions, zDEVD-FMK inhibited CASP1 probe cleavage with an IC_50_ of 15 μM, compared with 0.14 μM for Ac-YVAD-CMK, 1.7 μM for VX-765, and 0.04 μM for zVAD-FMK (Figure 3D,F). We performed an analogous analysis with CASP3 using its preferred substrate, Ac-DEVD-AMC. zDEVD-FMK inhibited CASP3 with an IC_50_ of 2 μM, whereas Ac-YVAD-CMK inhibited CASP3 with an IC_50_ of 0.54 μM. VX-765 exhibited little activity toward CASP3 (IC_50_ > >100 μM), while zVAD-FMK remained highly potent (IC_50_ = 0.099 μM) (Figure 3E,F), consistent with previous reports^39^. Thus, although CASP1 and CASP3 displayed the expected preferences for WEHD- and DEVD-based probes, respectively, these tetrapeptide preferences did not confer equivalent selectivity to the corresponding inhibitors.

We next asked whether inhibitor potency measured using fluorogenic probes reflected inhibition of endogenous protein substrate cleavage. Recombinant CASP1 was incubated with zDEVD-FMK, Ac-YVAD-CMK, VX-765, or zVAD-FMK before the addition of THP-1 macrophage lysates, and substrate cleavage was assessed by immunoblotting. Although zDEVD-FMK only weakly inhibited cleavage of the optimal CASP1 fluorogenic probe (IC_50_ = 15 μM), it inhibited CASP1-mediated processing of IL-18 and IL-1β at concentrations as low as 1 μM (Figure 1). Like zDEVD-FMK, Ac-YVAD-CMK and VX-765 partially inhibited IL-18 and IL-1β processing at 1 μM but failed to completely inhibit cytokine processing at 0.1 μM, whereas GSDMD cleavage required substantially higher inhibitor concentrations (Figure 3G). In contrast, zVAD-FMK efficiently inhibited processing of IL-18, IL-1β, and GSDMD across all concentrations tested (Figure 3G). These findings demonstrate that inhibition of fluorogenic probe cleavage does not accurately predict inhibition of protein substrates. Furthermore, DEVD- and YVAD-based inhibitors preferentially inhibited cytokine processing over GSDMD cleavage, providing additional evidence for substrate-dependent inhibition.

Because Ac-YVAD-CMK inhibited CASP3 fluorogenic probe cleavage more potently than zDEVD-FMK, we next asked whether this unexpected cross-reactivity also extended to endogenous protein substrates. Recombinant CASP3 was incubated with each inhibitor before the addition of THP-1 macrophage lysates. Consistent with the fluorogenic probe assay, zDEVD-FMK, Ac-YVAD-CMK, and zVAD-FMK inhibited cleavage of the endogenous CASP3 substrates PARP and GSDME, whereas VX-765 had no detectable effect (Figure 3H). Direct inhibition of CASP3 by Ac-YVAD-CMK was confirmed using purified recombinant IL-18, where cleavage was inhibited beginning at 0.16 μM (Figure 3I).

Taken together, these findings demonstrate that fluorogenic tetrapeptide probes are poor predictors of inhibitor activity toward endogenous protein substrates. Although DEVD- and YVAD-based inhibitors were developed to selectively target executioner and inflammatory caspases, respectively, both exhibit substantial protease cross-reactivity and substrate-dependent inhibition, highlighting the limitations of relying solely on tetrapeptide probe assays to define caspase inhibitor specificity.

### Low micromolar zDEVD-FMK preferentially inhibits cytokine processing during canonical inflammasome activation

Having established that zDEVD-FMK directly inhibits CASP1-mediated cytokine processing *in vitro*, we next asked whether this selectivity is maintained during canonical inflammasome activation in cells. THP-1 macrophages were pretreated with five-fold dilution series of zDEVD-FMK, Ac-YVAD-CMK, VX-765, or zVAD-FMK before stimulation with the NLRP3 agonist nigericin.

We first monitored pyroptotic pore formation by measuring Sytox Green uptake. Despite directly inhibiting CASP1 *in vitro*, zDEVD-FMK had no detectable effect on Sytox Green uptake at any concentration tested, similar to Ac-YVAD-CMK and zVAD-FMK (Figure 4A,B,D,E). Only VX-765 significantly reduced pore formation at 2 and 10 μM (Figure 4C,E). These findings suggested that inhibition of CASP1-mediated cytokine processing does not necessarily translate into inhibition of pyroptotic membrane permeabilization.

**Figure 4.**
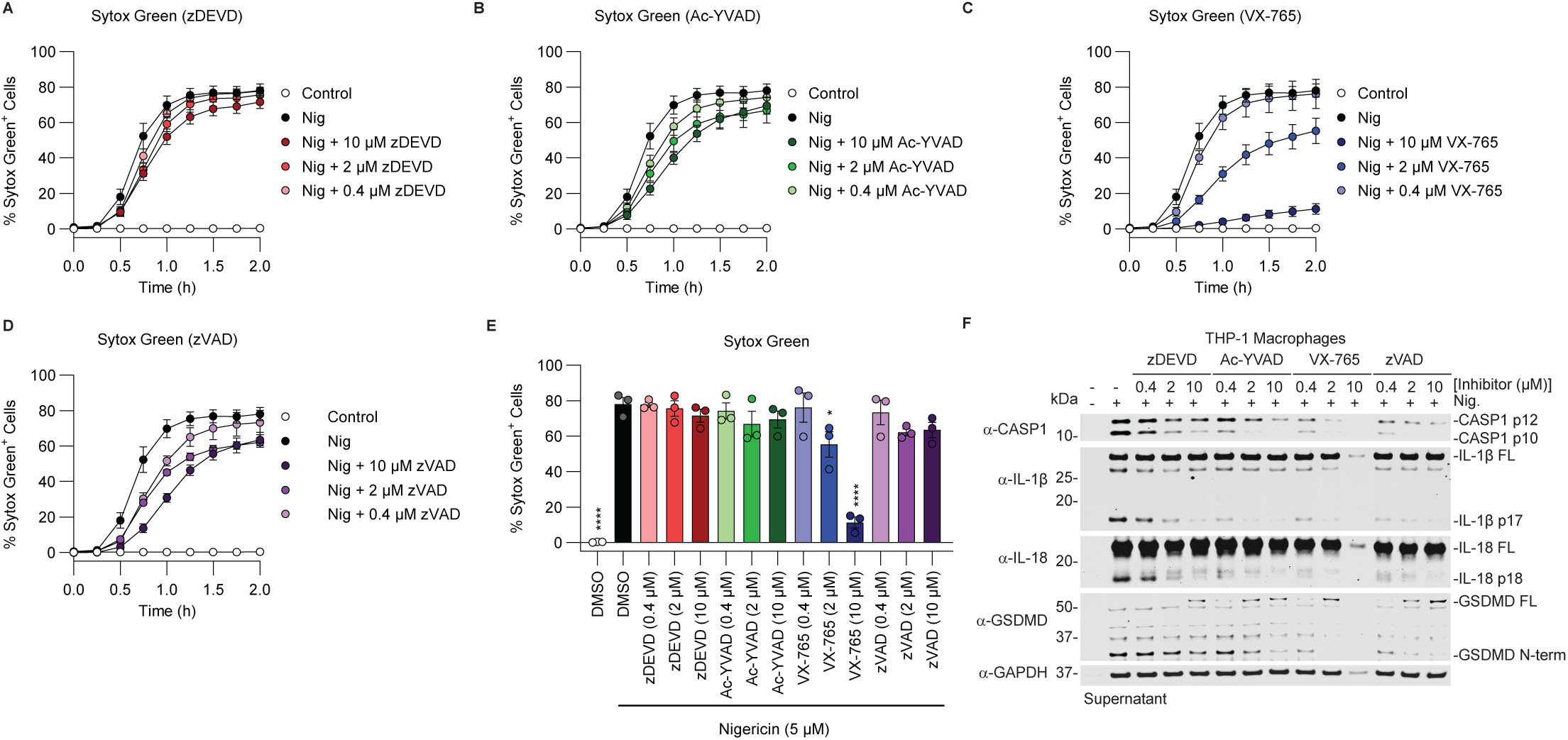
Low micromolar zDEVD-FMK preferentially inhibits cytokine processing during canonical inflammasome activation. (A-D) Time courses of Sytox Green uptake in PMA-differentiated THP-1 macrophages pretreated with the indicated concentrations of zDEVD-FMK (A), Ac-YVAD-CMK (B), VX-765 (C), or zVAD-FMK (D) before stimulation with 5 μM nigericin. (E) Quantification of Sytox Green uptake at 2 h after nigericin treatment. (F) Immunoblot analysis of CASP1 p10 generation and processing of IL-1β, IL-18, and GSDMD in culture supernatants from cells treated with the indicated inhibitors. Data are presented as mean ± SEM from three independent replicates and are representative of at least three independent experiments. ****P < 0.0001, ***P < 0.001, **P < 0.01, and *P < 0.05 by one-way ANOVA with multiple comparisons versus nigericin treatment alone at the 2 h time point.

We therefore asked whether individual CASP1 substrates differed in their sensitivity to pharmacologic inhibition. Culture supernatants collected 2 h after nigericin stimulation were analyzed by immunoblotting. zDEVD-FMK reduced, but did not completely abolish, generation of the CASP1 p10 fragment and processing of IL-1β and IL-18 across all concentrations tested (Figure 4F). In contrast, Ac-YVAD-CMK, VX-765, and zVAD-FMK nearly eliminated CASP1 processing and cytokine maturation at 2 and 10 μM and substantially reduced processing at 0.4 μM (Figure 4F).

Consistent with our biochemical assays, zDEVD-FMK had little effect on GSDMD cleavage despite reducing IL-1β and IL-18 processing (Figure 4F). Ac-YVAD-CMK modestly reduced GSDMD cleavage only at 10 μM, whereas VX-765 and zVAD-FMK inhibited GSDMD processing across all concentrations tested, with minimal residual cleavage at higher concentrations (Figure 4F). Together, these findings demonstrate that endogenous CASP1 substrates differ in their sensitivity to pharmacologic inhibition during canonical inflammasome activation. Specifically, zDEVD-FMK preferentially inhibits cytokine processing while sparing GSDMD cleavage, extending our biochemical observations to intact cells and establishing substrate-dependent inhibition as a feature of cellular inflammasome signaling.

### Low micromolar zDEVD-FMK inhibits fluorogenic CASP3/7 reporter activity but only modestly inhibits endogenous substrate cleavage during apoptosis

Because low micromolar concentrations of zDEVD-FMK inhibited CASP1-mediated cytokine processing during canonical inflammasome activation, we next asked whether these same concentrations effectively inhibit executioner caspase activity during apoptosis. THP-1 macrophages were pretreated with five-fold dilution series of zDEVD-FMK, Ac-YVAD-CMK, VX-765, or zVAD-FMK before stimulation with the apoptotic agonist staurosporine. Pore formation was monitored by Sytox Green uptake, executioner caspase activity was assessed using a fluorogenic CASP3/7 reporter containing a DEVD recognition sequence, and membrane lysis was quantified by LDH release.

Consistent with our observations during pyroptosis, zDEVD-FMK had no detectable effect on Sytox Green uptake at any concentration tested (Figure 5A,E). Ac-YVAD-CMK modestly reduced Sytox Green uptake at 10 μM, whereas VX-765 reduced uptake at 2 and 10 μM and zVAD-FMK reduced uptake only at 10 μM (Figure 5B-E). In contrast, zDEVD-FMK and zVAD-FMK nearly abolished CASP3/7 reporter activity at all concentrations tested, while Ac-YVAD-CMK and VX-765 significantly reduced reporter activity only at 10 μM (Figure 5F-J). Despite the marked inhibition of fluorogenic reporter cleavage, zDEVD-FMK did not reduce LDH release at any concentration tested. Ac-YVAD-CMK and VX-765 significantly reduced LDH release at 2 and 10 μM, whereas zVAD-FMK reduced LDH release only at 10 μM (Figure 5K).

**Figure 5.**
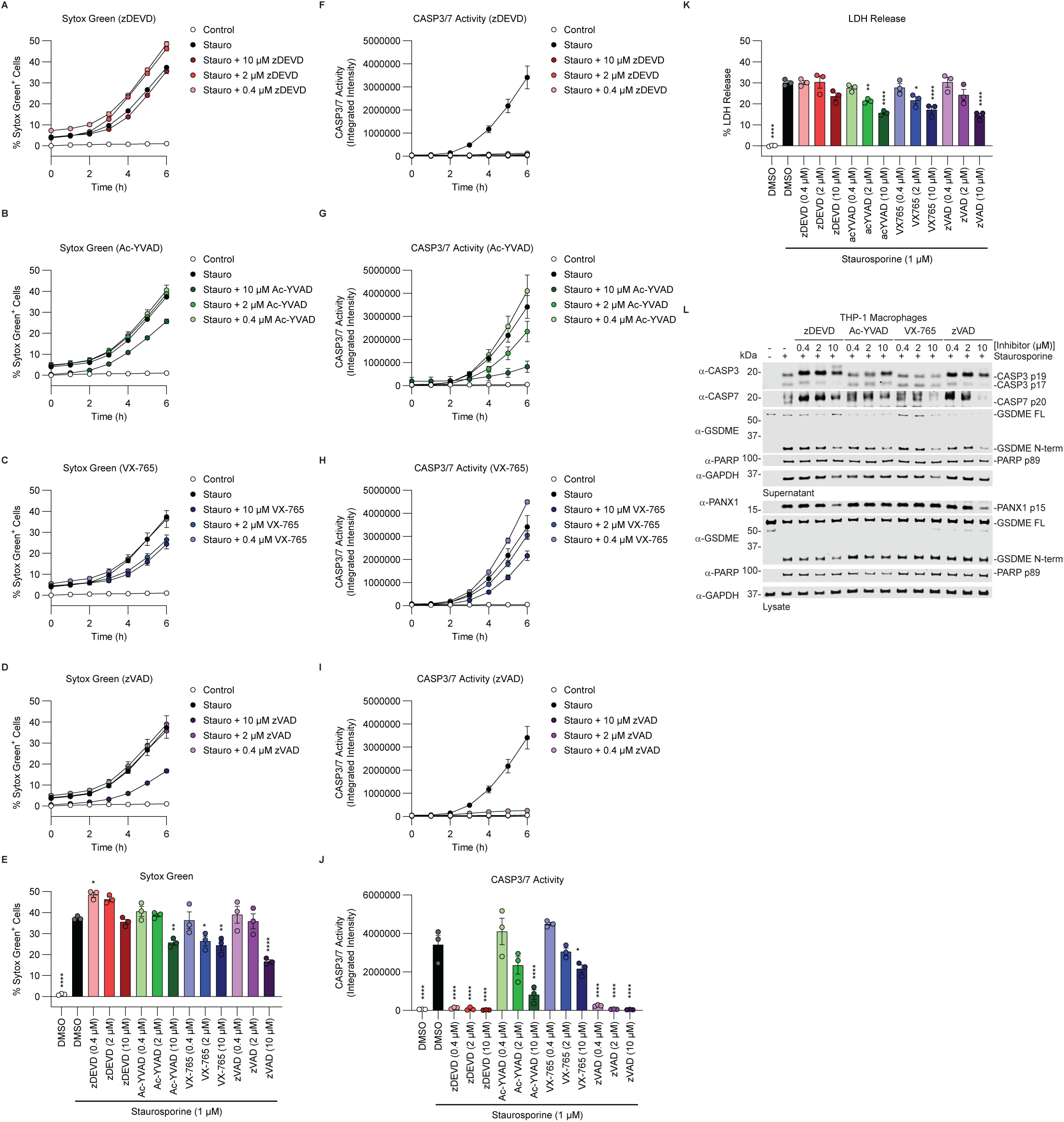
Low micromolar zDEVD-FMK inhibits fluorogenic CASP3/7 reporter activity but only modestly inhibits endogenous substrate cleavage during apoptosis. (A-D) Time courses of Sytox Green uptake in PMA-differentiated THP-1 macrophages pretreated with the indicated concentrations of zDEVD-FMK (A), Ac-YVAD-CMK (B), VX-765 (C), or zVAD-FMK (D) before stimulation with 1 μM staurosporine. (E) Quantification of Sytox Green uptake at 6 h after staurosporine treatment. (F-I) Time courses of CellEvent CASP3/7 reporter activity in cells treated with zDEVD-FMK (F), Ac-YVAD-CMK (G), VX-765 (H), or zVAD-FMK (I). (J) Quantification of CASP3/7 reporter activity at 6 h after staurosporine treatment. (K) LDH release at 6 h after staurosporine treatment. (L) Immunoblot analysis of CASP3 and CASP7 processing and cleavage of GSDME, PARP, and PANX1 in cell lysates and culture supernatants from cells treated with the indicated inhibitors. Data are presented as mean ± SEM from three independent replicates and are representative of at least three independent experiments. ****P < 0.0001, ***P < 0.001, **P < 0.01, and *P < 0.05 by one-way ANOVA with multiple comparisons versus staurosporine treatment alone at the 6 h time point for (E), (J), and (K).

We next asked whether inhibition of the fluorogenic reporter accurately reflected inhibition of endogenous CASP3/7 substrates. Cell lysates and culture supernatants were therefore analyzed for cleavage of GSDME, PARP, and PANX1 by immunoblotting. Surprisingly, despite nearly complete inhibition of the DEVD reporter, zDEVD-FMK and zVAD-FMK produced only modest reductions in cleavage of GSDME, PARP, and PANX1, even at 10 μM (Figure 5L). Likewise, Ac-YVAD-CMK and VX-765 had no detectable effect on endogenous substrate cleavage despite partially inhibiting fluorogenic reporter activity (Figure 5L).

Collectively, these findings demonstrate that inhibition of a DEVD fluorogenic reporter substantially overestimates inhibition of endogenous CASP3/7 substrates. Although zDEVD-FMK efficiently inhibits DEVD-based reporter cleavage and CASP1-mediated cytokine processing, it only modestly inhibits endogenous apoptotic substrate cleavage at the same concentrations. These results further demonstrate that inhibition of fluorogenic tetrapeptide probes does not faithfully predict inhibition of endogenous protein substrates.

### zDEVD-FMK cannot distinguish apoptotic and inflammatory caspase activity during LPS-induced non-canonical inflammasome activation

Because zDEVD-FMK inhibited CASP1-mediated cytokine processing during canonical inflammasome activation, we next asked whether it also suppresses inflammasome-dependent cytokine maturation during non-canonical inflammasome activation. This model additionally allowed us to determine whether zDEVD-FMK could distinguish inflammatory caspase activity from apoptotic executioner caspase activity as both pathways are activated concurrently.

Cytosolic lipopolysaccharide (LPS) activates the non-canonical inflammasome, leading to CASP4/5-mediated GSDMD cleavage, which can lead to downstream activation of the NLRP3 inflammasome and CASP1, and activation of CASP3 and CASP7^37,48^. THP-1 macrophages were pretreated with five-fold dilution series of zDEVD-FMK, Ac-YVAD-CMK, VX-765, or zVAD-FMK before transfection with 2.5 μg/mL LPS. Cell death was monitored by Sytox Green uptake and LDH release, executioner caspase activity was measured using a fluorogenic CASP3/7 reporter, and inflammasome-dependent cytokine signaling was assessed using HEK-Blue IL-1R reporter cells as we previously described and immunoblotting^28^.

Only zVAD-FMK significantly reduced Sytox Green uptake, and only at 10 μM, although Ac-YVAD-CMK showed a trend toward reduced uptake at the highest concentration tested (Figure 6A-E). In contrast, zDEVD-FMK and zVAD-FMK significantly reduced CASP3/7 reporter activity across all concentrations tested, whereas Ac-YVAD-CMK inhibited reporter activity at 2 and 10 μM and VX-765 only at 10 μM (Figure 6F-J). Despite having no detectable effect on Sytox Green uptake, zDEVD-FMK significantly reduced LDH release at 2 and 10 μM. VX-765 and zVAD-FMK reduced LDH release across all concentrations tested, whereas Ac-YVAD-CMK was effective only at 10 μM (Figure 6K).

**Figure 6.**
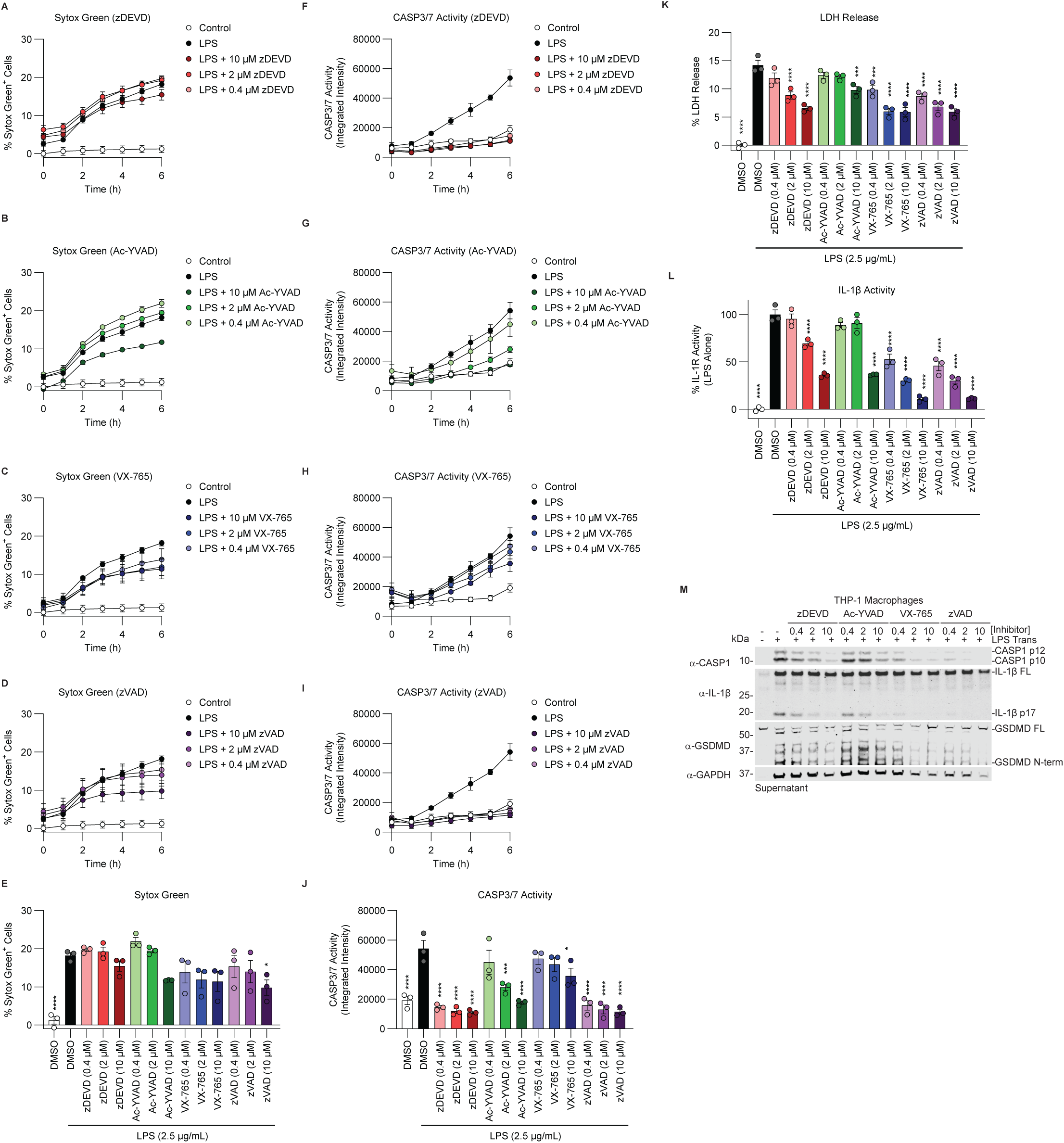
zDEVD-FMK cannot distinguish apoptotic and inflammatory caspase activity during LPS-induced non-canonical inflammasome activation. (A-D) Time courses of Sytox Green uptake in PMA-differentiated THP-1 macrophages pretreated with the indicated concentrations of zDEVD-FMK (A), Ac-YVAD-CMK (B), VX-765 (C), or zVAD-FMK (D) before transfection with 2.5 μg/mL LPS. (E) Quantification of Sytox Green uptake at 6 h after LPS transfection. (F-I) Time courses of CellEvent CASP3/7 reporter activity in cells treated with zDEVD-FMK (F), Ac-YVAD-CMK (G), VX-765 (H), or zVAD-FMK (I). (J) Quantification of CASP3/7 reporter activity at 6 h after LPS transfection. (K) LDH release at 6 h after LPS transfection. (L) IL-1α/β activity in culture supernatants measured using HEK-Blue IL-1R reporter cells. Reporter activity was normalized to untreated cells (0%) and LPS-transfected cells (100%). (M) Immunoblot analysis of CASP1 p10 generation and processing of IL-1β and GSDMD in culture supernatants from cells treated with the indicated inhibitors. Data are presented as mean ± SEM from three independent replicates and are representative of at least three independent experiments. ****P < 0.0001, ***P < 0.001, **P < 0.01, and *P < 0.05 by one-way ANOVA with multiple comparisons versus LPS transfection alone at the 6 h time point for (E), (J), (K), and (L).

We next determined whether zDEVD-FMK inhibited inflammasome-mediated cytokine maturation under these conditions. zDEVD-FMK significantly reduced IL-1R signaling at 2 and 10 μM, whereas Ac-YVAD-CMK reduced signaling only at 10 μM. VX-765 and zVAD-FMK inhibited IL-1R signaling across all concentrations tested (Figure 6L). Consistent with these findings, immunoblot analysis showed reduced generation of the CASP1 p10 fragment and IL-1β processing with all inhibitors, with VX-765 and zVAD-FMK producing nearly complete inhibition throughout the concentration range examined (Figure 6M). zDEVD-FMK and Ac-YVAD-CMK reduced GSDMD cleavage only at 10 μM, whereas VX-765 and zVAD-FMK inhibited GSDMD processing at all concentrations tested (Figure 6M).

Collectively, these findings demonstrate that zDEVD-FMK inhibits CASP1-mediated cytokine processing and fluorogenic CASP3/7 reporter activity over the same concentration range during LPS-induced non-canonical inflammasome activation. Thus, in a defined model of concurrent apoptotic and inflammatory caspase activation, DEVD-based inhibitors cannot distinguish executioner caspase activity from inflammasome signaling, highlighting the limitations of using tetrapeptide inhibitors to assign pathway-specific functions when multiple caspase pathways are engaged simultaneously.

### DEVD-based inhibitors cannot distinguish apoptotic and inflammatory caspase activity during *Salmonella* infection

We asked whether zDEVD-FMK could distinguish apoptotic executioner caspase activity from CASP1 activity during a physiologically relevant infection in which both pathways are activated concurrently. In THP-1 macrophages, *Salmonella enterica* serovar Typhimurium (*Stm*) activates CASP4 and CASP5, leading to activation of CASP3 and CASP7 while simultaneously triggering the canonical NLRC4 and NLRP3 inflammasomes (36). THP-1 macrophages were pretreated with five-fold dilution series of zDEVD-FMK, Ac-YVAD-CMK, VX-765, or zVAD-FMK (starting at 10 μM) before infection. Cell death was monitored by Sytox Green uptake, and executioner caspase activity was measured using the fluorogenic CASP3/7 reporter.

zDEVD-FMK, Ac-YVAD-CMK, and VX-765 each reduced Sytox Green uptake at 10 μM, whereas zVAD-FMK reduced Sytox Green uptake at both 2 and 10 μM (Figure 7A-E). zDEVD-FMK, Ac-YVAD-CMK, and zVAD-FMK significantly reduced CASP3/7 reporter activity at all concentrations tested, although none completely abolished reporter fluorescence (Figure 7F,G,I,J). VX-765 reduced reporter activity at 2 μM but unexpectedly showed a trend toward increased reporter activity at 10 μM (Figure 7H,J). Only Ac-YVAD-CMK and zVAD-FMK significantly reduced LDH release, and only at 10 μM (Figure 7K).

**Figure 7.**
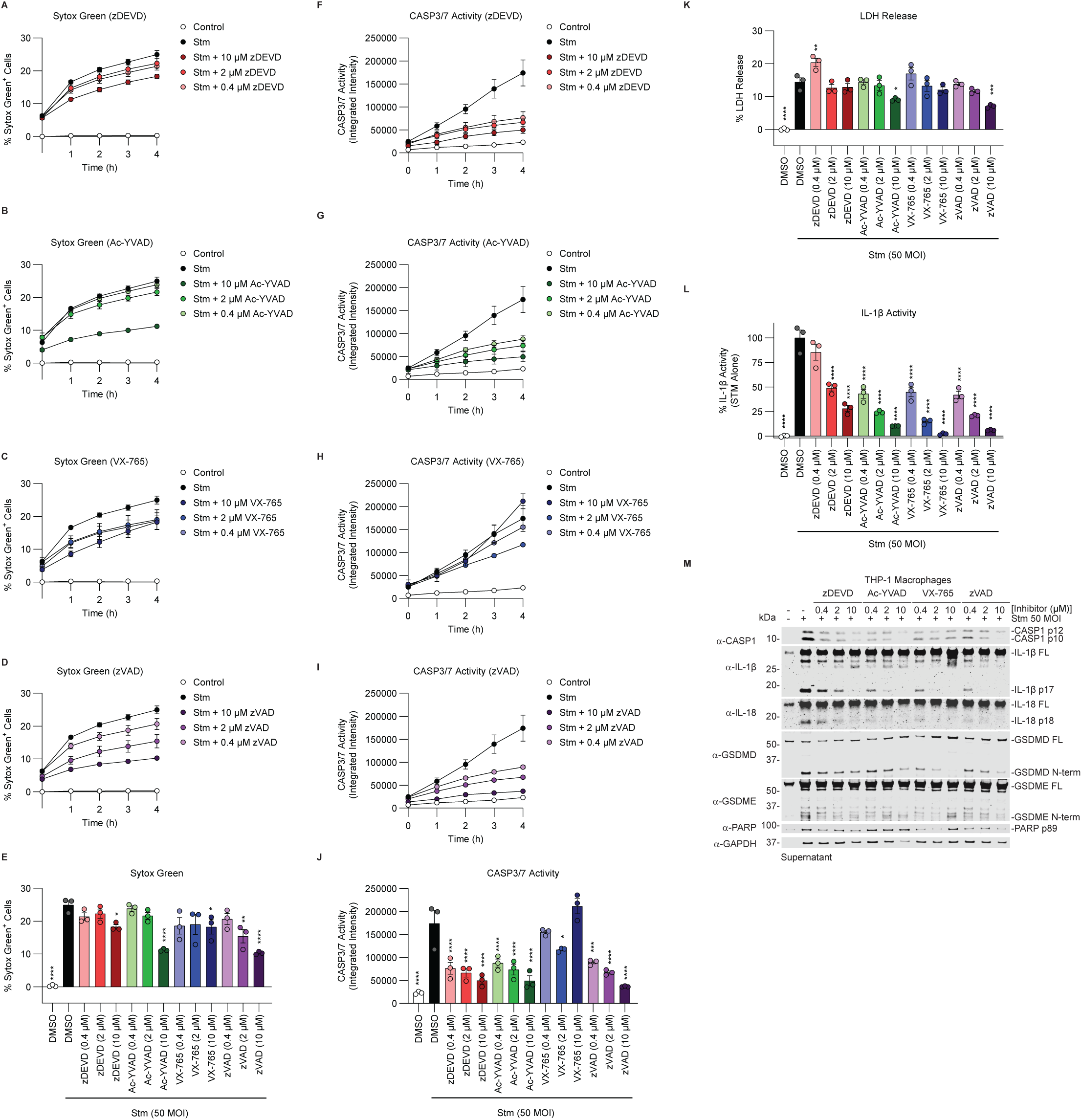
DEVD-based inhibitors cannot distinguish apoptotic and inflammatory caspase activity during *Salmonella* infection. (A-D) Time courses of Sytox Green uptake in PMA-differentiated THP-1 macrophages pretreated with zDEVD-FMK (A), Ac-YVAD-CMK (B), VX-765 (C), or zVAD-FMK (D) before infection with *Salmonella enterica* serovar Typhimurium. (E) Quantification of Sytox Green uptake at 4 h after *Salmonella* infection. (F-I) Time courses of CellEvent CASP3/7 reporter activity in cells treated with zDEVD-FMK (F), Ac-YVAD-CMK (G), VX-765 (H), or zVAD-FMK (I). (J) Quantification of CASP3/7 reporter activity at 4 h after *Salmonella* infection. (K) LDH release at 4 h after *Salmonella* infection. (L) IL-1α/β activity in culture supernatants measured using HEK-Blue IL-1R reporter cells. Reporter activity was normalized to untreated cells (0%) and *Salmonella*-infected cells (100%). (M) Immunoblot analysis of culture supernatants showing generation of the CASP1 p10 fragment, processing of IL-1β and IL-18, and cleavage of GSDMD, GSDME, and PARP in cells treated with the indicated inhibitors. Data are presented as mean ± SEM from three independent replicates and are representative of at least three independent experiments. ****P < 0.0001, ***P < 0.001, **P < 0.01, and *P < 0.05 by one-way ANOVA with multiple comparisons versus *Salmonella* infection alone at the 4 h time point for (E), (J), (K), and (L).

To assess inflammasome-dependent cytokine release, culture supernatants were analyzed using HEK-Blue IL-1R reporter cells. zDEVD-FMK reduced IL-1R signaling at 2 and 10 μM, whereas Ac-YVAD-CMK, VX-765, and zVAD-FMK significantly reduced IL-1R signaling at all concentrations tested (Figure 7L). Consistent with these findings, immunoblot analysis showed that all inhibitors reduced generation of the CASP1 p10 fragment and processing of IL-1β and IL-18 (Figure 7M). In contrast, zDEVD-FMK had no detectable effect on GSDMD cleavage, whereas Ac-YVAD-CMK, VX-765, and zVAD-FMK reduced GSDMD processing at 10 μM (Figure 7M). All inhibitors reduced GSDME cleavage except VX-765 at 10 μM, consistent with the increased CASP3/7 reporter activity observed under this condition (Figure 7H,J). zDEVD-FMK and zVAD-FMK modestly reduced PARP cleavage across the concentration range tested, Ac-YVAD-CMK had no detectable effect on PARP cleavage, and VX-765 reduced PARP cleavage at lower concentrations but not at 10 μM (Figure 7M).

Together, these findings extend our observations to a physiologically relevant bacterial infection model. During *Salmonella* infection, where apoptotic and inflammatory caspases are activated simultaneously, zDEVD-FMK inhibits CASP1-mediated cytokine processing while only modestly affecting endogenous apoptotic substrate cleavage. These results demonstrate that DEVD-based inhibitors cannot reliably distinguish executioner caspase activity from inflammasome signaling during infection and underscore the need for more selective caspase inhibitors to accurately define the contributions of these overlapping cell death pathways.

### Caspase inhibitors exhibit context-dependent substrate inhibition during canonical inflammasome activation in primary human macrophages

Having demonstrated that zDEVD-FMK preferentially inhibits CASP1-mediated cytokine processing over GSDMD cleavage in THP-1 macrophages, we next asked whether this substrate-dependent inhibition is conserved in primary human macrophages. To address this, we generated human monocyte-derived macrophages and pretreated them with VX765, zVAD-FMK, Ac-YVAD-CMK, or zDEVD-FMK at 2 or 10 μM before inducing canonical inflammasome activation with LPS and ATP. CASP1 substrate processing was assessed by immunoblotting of cell lysates and culture supernatants.

Unexpectedly, only VX765 inhibited IL-1β processing, whereas zVAD-FMK, Ac-YVAD-CMK, and zDEVD-FMK had no detectable effect at either concentration tested (Figure 8). In contrast, all four inhibitors suppressed cleavage of GSDMD and the apoptotic substrates GSDME and PARP at both concentrations (Figure 8). These findings contrast with our observations in THP-1 macrophages, where zDEVD-FMK preferentially inhibited cytokine processing while largely sparing GSDMD cleavage. Together, these results demonstrate that the substrate-dependent effects of caspase inhibitors vary between cellular models. Notably, zDEVD-FMK inhibits GSDMD cleavage without preventing IL-1β maturation in primary human macrophages, highlighting the importance of cellular context when evaluating caspase inhibitor specificity.

**Figure 8.**
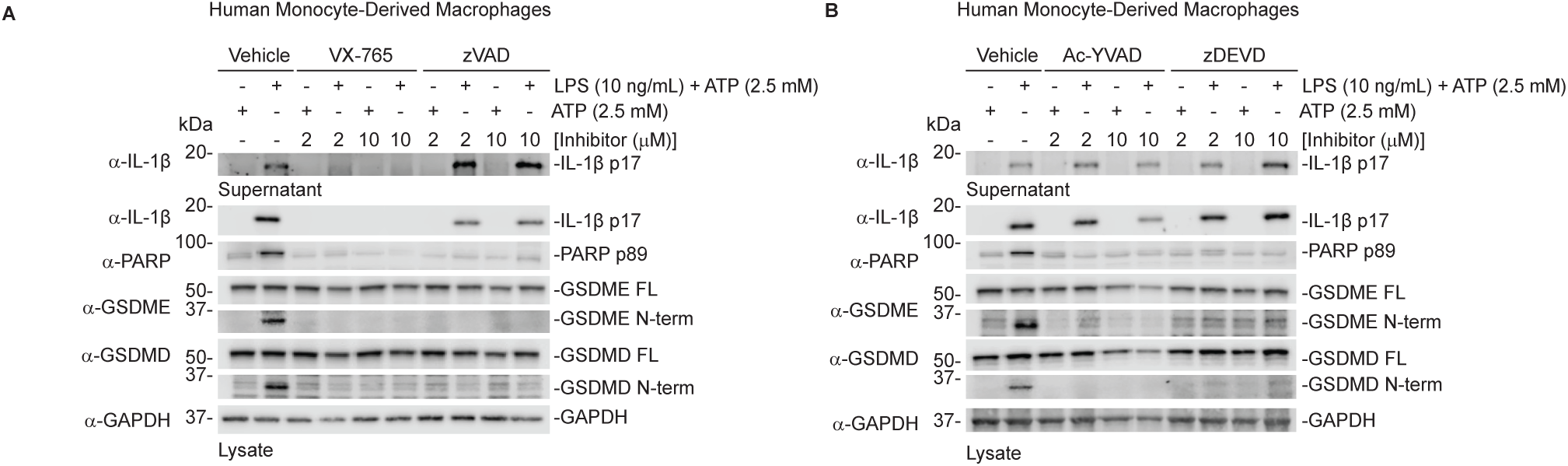
Caspase inhibitors exhibit context-dependent substrate inhibition during canonical inflammasome activation in primary human macrophages. (A,B) Immunoblot analysis of IL-1β processing and cleavage of GSDMD, GSDME, and PARP in culture supernatants and cell lysates from primary human monocyte-derived macrophages pretreated with VX-765, zVAD-FMK (A), or Ac-YVAD-CMK, zDEVD-FMK (B) (2 or 10 μM) before stimulation with LPS (10 ng/mL) and ATP (2.5 mM).

## Discussion

Peptide-based caspase inhibitors have been indispensable tools for studying programmed cell death and assigning biological functions to individual caspases. These inhibitors were developed from tetrapeptide substrate preferences identified using positional scanning libraries and validated using fluorogenic peptide substrates, with the underlying assumption that inhibition of peptide cleavage predicts inhibition of endogenous protein substrates. Here, we show that this assumption does not always hold. The widely used executioner caspase inhibitor zDEVD-FMK directly inhibited CASP1-mediated cytokine processing *in vitro* and in cells despite CASP1 exhibiting minimal activity toward the DEVD fluorogenic substrate. Conversely, the canonical CASP1 inhibitor Ac-YVAD-CMK directly inhibited CASP3 protein substrate cleavage. Together, these findings demonstrate that fluorogenic tetrapeptide substrates do not reliably predict inhibition of endogenous protein substrates and that peptide-based caspase inhibitors exhibit broader selectivity than is appreciated from conventional fluorogenic assays.

Beyond revealing unexpected protease cross-reactivity, our findings uncover an underappreciated feature of caspase pharmacology, that inhibitor potency depends not only on the protease being targeted but also on the substrate being measured and the biological context. Across multiple biochemical and cellular models, zDEVD-FMK consistently inhibited CASP1-mediated processing of IL-1β and IL-18 at concentrations that had little or no effect on GSDMD cleavage in THP-1 macrophages. Similar substrate-dependent differences were observed with Ac-YVAD-CMK and VX-765, indicating that this phenomenon is not unique to DEVD-based inhibitors. Likewise, during apoptosis, zDEVD-FMK nearly abolished cleavage of a DEVD fluorogenic reporter while only modestly reducing cleavage of endogenous CASP3 substrates, including PARP, GSDME, and PANX1^37,49^. Unexpectedly, in primary human monocyte-derived macrophages stimulated with LPS and ATP, zDEVD-FMK inhibited GSDMD cleavage without preventing IL-1β maturation, contrasting with its inhibitory profile in THP-1 macrophages. These findings demonstrate that the relative sensitivity of protein substrates to caspase inhibitors can vary substantially across experimental contexts. Thus, inhibition of one substrate cannot be assumed to reflect inhibition of other downstream functions of the same caspase.

The molecular basis for substrate-dependent inhibition remains unclear, but one reason could be that endogenous protein substrates and fluorogenic peptides engage caspases through fundamentally different molecular interactions. Fluorogenic substrates measure recognition of only the four amino acids immediately surrounding the scissile bond, whereas endogenous substrates present extended primary sequence, higher-order structure, and potential exosite interactions that contribute to substrate recognition and catalysis^29,30,47,50,51^. Differences in substrate affinity, accessibility, catalytic efficiency, or enzyme-substrate residence time may therefore influence apparent inhibitor potency. Cellular factors, including substrate abundance, subcellular localization, inhibitor availability, and the magnitude or kinetics of caspase activation, may further influence inhibition profiles. These considerations are particularly relevant to the differences observed between THP-1 and primary human macrophages. However, because these experiments also employed different inflammasome stimuli, additional studies will be required to distinguish cell-type-specific effects from differences in activation conditions. Nevertheless, our findings demonstrate that fluorogenic peptide assays alone are insufficient to establish inhibitor specificity and should be complemented by analyses of endogenous protein substrates in relevant cellular models.

Surprisingly, we also found that DEVD-based inhibitors preferentially inhibited CASP1 over the closely related inflammatory caspases CASP4 and CASP5. Tandem mass spectrometry identified covalent modification of the catalytic cysteine of CASP1 and CASP3 by zDEVD-FMK, whereas no corresponding adduct was detected for CASP4. In contrast, zVAD-FMK modified the catalytic cysteine of all three caspases, and preincubation with zVAD-FMK prevented detectable zDEVD-FMK modification of CASP1 and CASP3. These findings provide direct evidence that differential active-site engagement contributes to the distinct inhibitory profiles of inflammatory caspases. Although the absence of detectable CASP4 modification does not exclude weak or transient inhibitor interactions, the results are consistent with our biochemical observations. The molecular basis for this selectivity remains unresolved but may involve differences in active-site architecture, conformational dynamics, or inhibitor binding and reactivity that are not captured by tetrapeptide substrate preferences alone.

The differential inhibition of cytokine maturation and GSDMD cleavage has important biological implications. CASP1 coordinates inflammatory cytokine processing and GSDMD-mediated pore formation, two major outputs of canonical inflammasome activation. In THP-1 macrophages, zDEVD-FMK preferentially inhibited IL-1β and IL-18 processing while largely preserving GSDMD cleavage and membrane permeabilization. Conversely, in primary human macrophages, zDEVD-FMK inhibited GSDMD cleavage without detectably reducing IL-1β maturation. These observations suggest that distinct inflammasome outputs can exhibit different sensitivities to pharmacological inhibition and that the direction of this selectivity may vary with experimental context. Although these findings raise the possibility of selectively modulating inflammatory cytokine maturation and pyroptotic cell death, the mechanisms underlying these differences remain to be established. Moreover, inhibition of GSDMD cleavage does not necessarily establish selective inhibition of CASP1, particularly in settings where multiple inflammatory caspases may contribute to substrate processing.

Our findings also have important implications for studies investigating crosstalk between apoptosis and pyroptosis. During nigericin treatment, LPS transfection, and *Salmonella* infection, zDEVD-FMK inhibited CASP1-mediated cytokine processing over concentrations that also inhibited fluorogenic CASP3/7 reporter activity, despite only modestly inhibiting endogenous apoptotic substrate cleavage. Conversely, Ac-YVAD-CMK inhibited CASP3-mediated protein substrate cleavage despite being widely used as a CASP1-selective inhibitor. These observations demonstrate that DEVD- and YVAD-based inhibitors cannot reliably distinguish apoptotic executioner caspase activity from inflammasome signaling when both pathways are activated simultaneously. Importantly, our primary macrophage findings further indicate that inhibitor effects characterized in transformed cell lines cannot necessarily be extrapolated to primary cells. Consequently, assigning pathway-specific functions based solely on pharmacological inhibition may be misleading, particularly during infection and inflammatory conditions where multiple caspases are activated concurrently.

Collectively, our findings highlight three considerations that govern the interpretation of caspase inhibitor specificity: the protease being targeted, the endogenous substrate being measured, and the biological context in which inhibition occurs. Closely related caspases can exhibit markedly different sensitivities to the same inhibitor, while distinct substrates of a single caspase can differ substantially in their sensitivity to inhibition. Moreover, these inhibitory profiles can vary across cellular models and activation conditions. Together, these observations demonstrate that caspase inhibitor specificity cannot be established solely through fluorogenic tetrapeptide assays or inhibition of a single downstream substrate. Instead, inhibitor activity should be evaluated using multiple endogenous protein substrates in biologically relevant models, ideally alongside orthogonal genetic approaches. These findings have broad implications for interpreting studies that rely on peptide-based caspase inhibitors and underscore the need for next-generation inhibitors with improved protease selectivity and more predictable effects on downstream substrate processing.

## Methods

### Antibodies and reagents

Antibodies used include: GSDMD Rabbit polyclonal Ab (Novus Biologicals, NBP2–33422), GAPDH Rabbit monoclonal Ab (Cell Signaling Tech, 14C10), CASP1 p12/10 Rabbit monoclonal Ab (Abcam, ab179515), CASP3 (Cell Signaling Tech 9664S), CASP7 (Cell Signaling Tech, 8438S), GSDME (Abcam, ab215191), hIL-1β Goat Polyclonal Ab (R&D systems, AF-201-NA), hIL-18 Goat Ab (R&D systems, af2548), Pannexin-1 (D9M1C) (Cell Signaling Tech, 91137), PARP (Cell Signaling Tech, 9542S). IRDye 800CW anti-rabbit (LICORbio, 925–32211), IRDye 800CW anti-mouse (LICORbio, 925–32210), IRDye 800CW anti-goat (LICORbio, 926-32214), IRDye 680CW anti-rabbit (LICORbio, 925–68073), IRDye 680CW anti-mouse (LICORbio, 925–68072), IRDye 800CW anti-goat (LICORbio, 926-68074).

Other reagents used include: Ac-DEVD-AMC (Fisher Scientific, ALX-260-031-M005), zDEVD-FMK (R&D Systems, FMK004), Ac-DEVD-CMK (MedChem Express, HY-P0034), Ac-DEVD-CHO (MedChem Express, HY-P1001), Ac-WEHD-AMC (Fisher Scientific, ALX-260-057-M005), Ac-YVAD-CMK (Invivogen, inh-yvad), zYVAD-FMK (MedChem Express, HY-P1009-1mg), VX-765 (Fisher Scientific, 50-101-3604), zVAD-FMK (Fisher Scientific, ALX-260-020-M005), Nigericin (sodium salt) (Cayman Chemical, 11437), Staurosporine (MedChemExpress, HY-15141), Caspase-1 (human), (recombinant) (Enzo Life Sciences, BML-SE168-5000), Caspase-3 (human), (recombinant) (Enzo Life Sciences, BML-SE169-5000), Caspase-4 (human), (recombinant) (Enzo Life Sciences, BML-SE176-5000), Caspase-5 (human), (recombinant) (Enzo Life Sciences, BML-SE171-5000), Caspase-7 (human), (recombinant) (Enzo Life Sciences, BML-SE177-5000), SimplyBlue SafeStain (Thermo Fischer Scientific, LC6065), phorbol 12-myristate 12-acetate (PMA) (Promega, V1171), Sytox Green (Invitrogen, S7020), CellEvent Caspase-3/7 Detection Reagent (Invitrogen, C10423), Pan Monocyte Isolation Kit (Miltenyi Biotec, #130-096-537), M-CSF (PeproTech, #315-02), GlutaMAX (Thermo Fisher Scientific, #51985034), LPS (MilliporeSigma, #L2880, ATP (Fisher Scientific, #AAJ6112506), *Escherichia coli* Rosetta (DE3) pLysS cells (Neta/Sigma Aldrich, 71-397-3), kanamycin (Sigma-Aldrich, K1377-5G), and chloramphenicol (Sigma-Aldrich, C0378-25G), isopropyl β-D-1-thiogalactopyranoside (IPTG) (Sigma-Aldrich, I6758-10G), HisTrap HP affinity column (Neta Scientific, 29051021), TEV protease (GenScript, Z03030), Amicon Ultra-15 centrifugal filter unit with a 10-kDa molecular weight cutoff (Fisher Scientific, UFC901024), MS grade trypsin/Lys-C mix (Promega, catalog no. V5072), Empore™ C_18_ extraction disks (cat. no. 66883-U).

### Expression and purification of recombinant human pro-IL-18

The pET28_human-pro-IL18 plasmid (Addgene #183390) was transformed into chemically competent *Escherichia coli* Rosetta (DE3) pLysS cells (Neta/Sigma Aldrich, 71-397-3). Transformants were selected on agar plates containing kanamycin (50 µg/mL) (Sigma-Aldrich, K1377-5G) and chloramphenicol (25 µg/mL)(Sigma-Aldrich, C0378-25G). A single colony was inoculated into 5 mL of 2×YT medium supplemented with both antibiotics and grown overnight at 37 °C with shaking at 225 rpm.

The overnight culture was used to inoculate 1 L of 2×YT medium containing kanamycin (50 µg/mL) and chloramphenicol (25 µg/mL) in a 2-L baffled flask. Cells were grown at 37 °C with shaking at 225 rpm until the optical density at 600 nm (OD₆₀₀) reached 0.6–0.8. Protein expression was induced with 0.25 mM isopropyl β-D-1-thiogalactopyranoside (IPTG) (Sigma-Aldrich, I6758-10G) for 4 h at 37 °C with shaking at 200 rpm. Cells were harvested by centrifugation at 5,000 × *g* for 15 min at 4 °C, and pellets were stored at −20 °C until purification.

Bacterial pellets were resuspended in 30–40 mL of lysis buffer (25 mM HEPES, pH 7.4, 150 mM NaCl, and 10 mM imidazole) and lysed by sonication using a Q125 sonicator (Qsonica) equipped with a ½-inch probe at 100% amplitude. Sonication was performed using 10-s pulses separated by 59-s cooling intervals for a cumulative sonication time of 5 min (34.5 min total processing time), with samples maintained in an ice-water bath. Lysates were clarified by centrifugation at 30,000 × *g* for 45 min at 4 °C and filtered through 0.45-µm syringe filters before loading onto a 1-mL HisTrap HP affinity column (Neta Scientific, 29051021) equilibrated with lysis buffer.

The column was washed with 10 mL of wash buffer (25 mM HEPES, pH 7.4, 400 mM NaCl, and 25 mM imidazole). His-tagged protein was eluted using an ÄKTA pure FPLC system with a linear imidazole gradient from 25 to 200 mM over 60 min at a flow rate of 1 mL/min. Elution fractions were analyzed by SDS-PAGE and SimplyBlue SafeStain staining. Fractions containing pro-IL-18 were pooled and dialyzed overnight at 4 °C against 25 mM HEPES (pH 7.4), 150 mM NaCl, and 1 mM DTT in the presence of TEV protease (GenScript, catalog no. Z03030; 150 IU/mL) to remove the His-tag.

TEV-mediated cleavage was verified by SDS-PAGE and SimplyBlue SafeStain staining. The protein was then applied to a second 1-mL HisTrap HP column equilibrated with 25 mM HEPES (pH 7.4) and 150 mM NaCl to remove His-tagged TEV protease and the cleaved His-tag. The flow-through containing untagged pro-IL-18 was collected and concentrated to approximately 1 mg/mL using an Amicon Ultra-15 centrifugal filter unit with a 10-kDa molecular weight cutoff (Fisher Scientific, UFC901024). Glycerol was added to a final concentration of 8% (v/v), and the purified protein was aliquoted, snap-frozen in liquid nitrogen, and stored at −80 °C.

### In vitro substrate cleavage assays

All *in vitro* assays using recombinant caspases were performed in 20 μL reactions containing caspase assay buffer (20 mM PIPES, 100 mM NaCl, 10 mM DTT, 1 mM EDTA, 0.1% CHAPS, 10% sucrose, pH 7). We used 1 activity unit/μL of each indicated caspase. For recombinant IL-18 cleavage assays, a final concentration of 50 μg/mL in caspase buffer was used. For THP-1 lysate experiments, 1.6 × 10^7^ THP-1 macrophages were differentiated in a 15cm dish in 50 ng/mL phorbol 12-myristate 12-acetate (PMA) (Promega, V1171). After 24 h, cells were harvested and lysed by sonication in 10 s pulses for 60 s at 30% amplitude in 10 mL PBS and centrifuged at 12,000 g to remove cell debris. Inhibitors and caspases were preincubated for 5 minutes before the addition of lysates diluted to a final dilution of 6x in caspase assay buffer. Reactions were incubated at 1 h at 37 °C. Samples were mixed 1:1 with 2× Licor Protein Sample Loading Buffer (Neta Scientific, 928-40004), boiled at 95 °C for 10 min, and then analyzed by SDS-PAGE and immunoblotting for THP-1 lysate experiments, or gels were washed 3x in tap water and stained with SimplyBlue SafeStain for recombinant IL-18 experiments.

### Caspase-Inhibitor Preparation for Mass Spectrometry

Caspases were incubated at 1 activity unit/μl in caspase buffer with indicated inhibitors for 60 minutes at room temperature. Where indicated caspases were first incubated with zVAD-FMK for 30 minutes followed by the addition of zDEVD-FMK for an additional 30 minutes. Subsequently, freshly prepared iodoacetamide (IA) was added to a final concentration of 20 mM, and samples were incubated at room temperature in the dark for 10 min to allow alkylation of cysteine residues. Proteins were digested by adding MS grade trypsin/Lys-C mix (Promega, catalog no. V5072) at an enzyme to protein ratio of 1:100 (w/w), followed by incubation at 37 °C overnight with shaking.

Digested peptides were desalted using C_18_ stage tips packed with Empore™ C_18_ extraction disks (cat. no. 66883-U) following a standard protocol. Stage tips were conditioned with methanol and 50% (v/v) acetonitrile containing 0.1% (v/v) formic acid, equilibrated with 0.1% (v/v) formic acid, and loaded with peptides reconstituted in 3% (v/v) acetonitrile and 0.1% (v/v) formic acid. After washing with 0.1% (v/v) formic acid, peptides were eluted using 50% (v/v) acetonitrile containing 0.1% (v/v) formic acid, vacuum-dried, and subsequently submitted for LC-MS/MS analysis.

### QTOF LC-MS/MS Analysis

LC-MS/MS analyses were conducted on an Agilent 1290 Infinity II UHPLC system coupled with an Agilent 6545XT AdvanceBio LC/Q-TOF system equipped with an Agilent Dual Jet Stream ESI source. LC separation was obtained with an Agilent AdvanceBio Peptide Mapping column (2.1 × 150 mm, 2.7 µm). Peptides were separated over a gradient of 1-95% acetonitrile in water (both containing 0.1% formic acid) over 45 minutes with a flow rate of 0.4 ml/min. MS and MS/MS data were acquired using the Q-TOF with the following parameters:

|  |  |
|---|---|
| Drying Gas | 11 L/min |
| Drying Gas Temperature | 325 °C |
| Sheath Gas Flow | 10 L/min |
| Sheath Gas Temperature | 325 °C |
| Nebulizer Pressure | 35 psi |
| Capillary Voltage | 4,000 V |
| Nozzle Voltage | 0 V |
| Fragmentor Voltage | 175 V |
| Acquisition Mode | AutoMS2 |

### LC–MS/MS database searching and inhibitor-adduct identification

Raw LC–MS/MS data were converted to mzML format and searched using FragPipe v23.1 with MSFragger v4.4.1. Spectra were searched against custom FASTA databases containing the mature subunits of the recombinant caspases analyzed, supplemented with common contaminant proteins and reversed decoy sequences. For the caspase-1/-3 experiments, the database contained mature caspase-1 and caspase-3 subunits; the caspase-4 analysis was performed using a database additionally containing mature caspase-4 and caspase-5 sequences.

Searches were performed using strict tryptic specificity together with Lys-C specificity, allowing up to three missed tryptic cleavages and two missed Lys-C cleavages. Peptides of 6–50 amino acids and 500–5000 Da were considered, with up to three variable modifications per peptide. The precursor true-mass tolerance was 20 ppm, isotope errors of 0, +1, or +2 were allowed, and b- and y-type fragment ions were considered. The optimized fragment-mass tolerance was 20 ppm.

Methionine oxidation (+15.9949 Da), protein N-terminal acetylation (+42.0106 Da), and cysteine carbamidomethylation (+57.02146 Da) were included as variable modifications. Carbamidomethylation was treated as variable rather than fixed to permit mutually exclusive assignment of alkylated and inhibitor-modified cysteine residues. Candidate inhibitor-derived modifications were searched using selected mass offsets of +415.17435, +433.18490, and +447.20056 Da for zVAD-derived species and +616.23810, +634.24860, and +648.26430 Da for zDEVD-derived species. The +433.1849- and +648.2643-Da species were also included explicitly as variable modifications.

To facilitate identification of covalently modified peptides and potentially labile inhibitor-derived fragment ions, MSFragger was operated in labile-search mode with delta-mass localization enabled. Remainder masses of 214.09536 and 242.12670 Da were included for zVAD- and zDEVD-derived fragmentation, respectively. For the initial analysis, inhibitor-derived delta masses were permitted on C, S, T, Y, K, and R to assess potential alternative sites of modification; subsequently the modifications were restricted to cysteine.

Peptide-spectrum matches were rescored using MSBooster v1.3.17 and Percolator v3.7.1 using a target-decoy strategy. Modification-site localization was further evaluated using PTMProphet v7.3.0 with a 10-ppm fragmention tolerance and a minimum localization probability of 0.5. Protein inference was performed using ProteinProphet, and final protein-level filtering was performed with Philosopher v5.1.2 using a picked target-decoy strategy and a 1% protein false-discovery rate.

Candidate inhibitor-modified active-site peptides were subsequently evaluated manually. Assignments were required to be consistent with the expected precursor mass shift and to localize the modification to the catalytic cysteine on the basis of fragment-ion evidence. Where multiple inhibitor-derived mass states were observed, spectra were inspected for diagnostic inhibitor fragments and for peptide fragment ions retaining the corresponding mass shift. Absence of labeling was assigned only when the corresponding catalytic-site peptide was confidently detected in the same sample without an inhibitor-derived modification.

### In Vitro Substrate Kinetics

All fluorogenic substrates were resuspended in DMSO at 40 mM stock concentrations and diluted to the indicated concentrations in caspase assay buffer. All probe experiments were completed at a 20 μL final volume, with 1 activity units per μL of each recombinant caspase. Relative fluorescence units were measured every 5 min for 30 minutes using a Cytation 5 plate reader (Biotek). Background fluorescence was subtracted from samples incubated with caspases. At each concentration of substrate, simple linear regressions (*y* = *mx* + *b*) were performed using GraphPad Prism to determine the rate. Rates were plotted against substrate concentration for n = 3 separate runs and fit to a Michaelis−Menten curve (*Y* = V_max_ × *X*/(K_M_ + *X*)) on Prism. For V_max_, K_M_, and k_cat_/K_M_ calculations, curves were generated for each individual experimental replicate, and V_max_, K_M_, and k_cat_ values were recorded for each replicate. k_cat_ was determined as the V_max_ value with the total enzyme concentration (ET) set to 1 for all caspases. The mean and standard error of the mean were calculated by compiling the values for each individual run.

### In Vitro Inhibition Assays

To determine the inhibition of maximal enzyme activity, each caspase was incubated with its optimal probe substrate at a saturating concentration. Inhibition of maximal caspase activity was determined with 200 μM Ac-WEHD-AMC for caspase-1 and 100 μM Ac-DEVD-AMC for caspase-3. Rates were determined as described in the *In Vitro Substrate Kinetics* section. Rates for n = 3 independent runs were plotted against inhibitor concentration. Percent caspase activity was determined by normalizing each independent run, with 0% activity set at 0 and maximum activity set as the sample with no inhibitor added. Replicates were combined, and inhibition curves were calculated for the aggregate data using an [inhibitor] vs normalized response curve on GraphPad Prism (*Y* = 100/ (1 + *X*/IC50)).

### Cell lines and culture methods

All cells were grown at 37 °C in a 5% CO_2_ atmosphere incubator. THP-1 cells were acquired from the Abbott lab (33). THP-1 cells were cultured in Roswell Park Memorial Institute (RPMI) medium 1640 with L-glutamine and 10% fetal bovine serum (FBS). Cell lines were negative for mycoplasma contamination.

### Testing Inhibitors in Cells

THP-1 cells were resuspended in RPMI medium containing 50 ng/mL phorbol 12-myristate 12-acetate (PMA) and plated in black, clear-bottom 96-well plates at a density of 8 × 10^4^ cells/well. After 24 h, the media was then replaced with serum-free RPMI (0.1 mL/well) supplemented with either CellEvent Caspase-3/7 Detection Reagent (final concentration: 10 µM (1:200 dilution)) or Sytox Green (final concentration: 0.5 µM) as indicated. Cells were treated with zDEVD-FMK, Ac-YVAD-CMK, VX-765 or zVAD-FMK 30 min before the cell death trigger was added at the indicated concentration. All inhibitors were resuspended in DMSO at 40 mM and diluted to the indicated concentration. DMSO concentration for all cell-based experiments was below 0.05%. Cells were incubated in the trigger for the indicated time points, and analyzed using IncuCyte live imaging. Supernatants and lysates were harvested and pooled across replicates, and proteins were prepared as described below. Nigericin (Neta Scientific, 11437) was resuspended in ethanol at 5 mM and added at a final concentration of 5 μM, staurosporine (MedChemExpress, HY-15141) was resuspended in DMSO at 1 mM and added at a final concentration of 1 μM, LPS (Cell Signaling Technology, 14011S) was resuspended in water at 5 mg/mL and mixed with Fugene HD (Promega, E2312) for at least 15 minutes in serum free media to a final concentration of 2.5 μg/mL LPS and 0.5% Fugene HD.

### IncuCyte CASP3/7 activity and Sytox Green uptake assays

The kinetics of CASP3/7 activity and Sytox Green uptake were measured in cells using the IncuCyte <u>S3</u> Live cell imaging system (Sartorius). A series of images were collected. The number of Sytox Green positive cells or CASP3/7 integrated intensity in each image was determined using the IncuCyte S3 software with 2 different areas/well. Cell death was quantified relative to 0.1% Triton-X treated macrophages.

### LDH assays

Supernatants were harvested and analyzed using the Cyquant LDH Cytotoxicity Assay (Thermo Scientific, C20301) according to the manufacturer’s protocol. LDH activity was quantified relative to a lysis control where cells were lysed using 0.1% Triton-X.

### Protein preparation for western blotting

For supernatant samples, proteins were precipitated using methanol-chloroform extraction. Methanol was mixed with supernatant samples at a 1:1 ratio and chloroform was added at 0.3:1. Samples were spun at 20,000g for 10 minutes and the aqueous layer was discarded. Proteins were then washed with methanol at a minimum 2:1 ratio and spun at 20,000g for 10 minutes. Methanol wash was removed and samples were dried at 65° C. Samples were resuspended in 2× Licor Protein Sample Loading Buffer (Neta Scientific, 928-40004) and boiled for 10 minutes at 95° C. For lysate samples, cells were lysed directly in 2× Licor Protein Sample Loading Buffer, sonicated for 10 s pulses for 60 s at 30% amplitude and boiled for 10 minutes at 95° C.

### Western blotting

Protein samples were run on Nupage 4–12% Bis-Tris Midi Protein Gel (Thermo Scientific, WG1403BOX) or Bolt™ Bis-Tris Plus Mini Protein Gels, 4-12%, WedgeWell™ format gel (Thermo Scientific, NW04127BOX) at 175 volts. Proteins were transferred to 0.45 µm nitrocellulose membranes (BioRad,1704271) at 25 volts for 7 minutes using the Bio-Rad Transblot Turbo System. Membranes were blocked using blocking buffer (3% Bovine Serum Albumin (Research Products International, A30075) in TBS buffer with 0.1% Tween and 0.05% Sodium Azide (VWR, BDH7465)) for 1 hour and stained with primary antibodies at 1:1000 concentration in blocking buffer overnight at 4° C. Membranes were washed 3x in TBS buffer with 0.1% Tween, followed by incubation with Donkey Anti-Mouse/Rabbit/Goat IgG Polyclonal Antibody for 1 hour at room temperature. Membranes were then washed 3x and imaged on the Odyssey M Imaging System (LICORbio). Images were analyzed with Empiria studio version 3.2 (LICORbio), and brightness, contrast, and tone parameters were adjusted uniformly in Adobe Photoshop.

### Salmonella infection of human macrophages

30 minutes prior to infection, indicated inhibitors were added to cells. *Salmonella* Typhimurium SL1344 was streaked to isolation from frozen stock on LB + Streptomycin (100 µg/mL) agar plates. To induce SPI-1, single colonies were inoculated in LB + Streptomycin broth overnight (∼18h). The overnight culture was back diluted 40-fold into LB + Streptomycin broth containing 300 mM NaCl. After three hours bacteria were washed 2x in sterile PBS and cell density was determined by quantifying OD_600_. MOI was prepared according to OD and bacterial cells were resuspended in PBS. 10µL suspension of bacteria was added to macrophages in triplicate in 96-well plates. Plates were spun for 10 min at 400g to allow infection and then placed in the 37°C in a 5% CO_2_ atmosphere incubator for 1 h. Media was then removed and replaced with fresh RPMI without FBS containing gentamicin (100 µg/mL), indicated inhibitors, and indicated fluorescent dyes and monitored using IncuCyte live cell imaging for 4 h.

### HEK blue IL-1α/β reporter assay

HEK-Blue IL-1β (Invivogen) reporter cells were seeded in 384 well plates at 1.5 × 10^4^ cells/well in DMEM + 10% FBS to a total volume of 20 µL. 5 µL of supernatant from treated THP-1 cells were added, and the samples were incubated overnight at 37°C in a 5% CO_2_ atmosphere incubator. 10 µL of supernatant was mixed with 90 µL of Quanti-Blue solution and incubated at room temperature for 10-30 minutes in the dark before reading absorbance at 620 nm using a Cytation 5 plate reader (Bio Tek). 0% IL-R activity is defined by a media only control while 100% activity is defined by the cell death trigger-treated sample with no inhibitor.

### Primary Cell Inhibitor Experiments

Primary human monocyte-derived macrophages (hMDMs) were generated from fresh human blood samples obtained from healthy donors through the New York Blood Center (NYBC). Peripheral blood mononuclear cells (PBMCs) were isolated by Ficoll density gradient centrifugation, and monocytes from 10 million PBMCs were isolated using the Pan Monocyte Isolation Kit (Miltenyi Biotec, #130-096-537) to determine monocyte population. Cells were then seeded in 6-well plates at a density of 950,000 monocytes/well and cultured in complete macrophage medium consisting of DMEM-GlutaMAX supplemented with 10% FBS, 5% human AB serum, 1% HEPES, and 1% penicillin/streptomycin, along with 10 ng/mL M-CSF (PeproTech, #315-02). On day 3, cells were washed and replenished with fresh M-CSF-supplemented medium to omit the non-adherent cells. Monocytes were allowed to fully differentiate into macrophages for 6-7 days *in vitro* (DIV). For inhibitor experiments, fully differentiated macrophages were switched to serum-free Opti-MEM Reduced Serum Medium supplemented with GlutaMAX (Thermo Fisher Scientific, #51985034) 1 h before treatment. Inhibitors were added directly to the wells at final concentrations of 2 or 10 µM, as indicated, and cells were preincubated for 45 min. DMSO was added to control wells as the vehicle control. Subsequently, cells were stimulated with LPS (10 ng/mL; MilliporeSigma, #L2880) or diH₂O as a control for 4 h. ATP (Fisher Scientific, #AAJ6112506) was then diluted in Opti-MEM to 5 mM and added during a half-medium exchange, resulting in a final ATP concentration of 2.5 mM in the wells. Following 1 h of ATP stimulation, lysates and supernatants were collected.

### Statistics

Data and statistical analysis were performed using GraphPad Prism 10 software. For each time course experiment, statistical analyzes were performed on the final timepoint. Statistical analysis was determined using One-way ANOVA with multiple comparisons tests where the cell death trigger-treated sample with no inhibitor is set as the control group.

## Author contributions CRediT

Conceptualization: CYT, CMB, Methodology: CYT, CMB, MD, PJG, JCJH, GMB, Investigation: CMB, JDL, KB, HTG, BMD, MD, JCJH, EM, Visualization: CYT, CMB, Funding acquisition: CYT, CMB, PJG Project administration: CYT, CMB, Data curation: CYT, CMB, GMB, Formal Analysis: CYT, CMB, JDL, KB, HTG, BMD, JCJH, GMB, Resources: CYT, Supervision: CYT, Writing – original draft: CYT, CMB, Writing – review & editing: CYT, CMB, JDL.

## Acknowledgements

This work was supported by an NIGMS Maximizing Investigator’s Research Award (Grant# 1R35GM155239-01 to CYT), the Penn Provost Postdoctoral Fellowship to CMB, and Burroughs Wellcome Fund (Grant# 1054907 to CMB), NIH/NIDA National Institute on Drug Abuse (R33DA05850103, R01DA062277 to PJG), the National Institute of Allergy and Infectious Diseases (F31AI203627 to MD), Schlumberger Faculty for the Future Fellowship (MD).

